# Paip2 associates with PABPC1 on mRNA, and may facilitate PABPC1 dissociation from mRNA upon deadenylation

**DOI:** 10.1101/2021.02.07.430161

**Authors:** Jingwei Xie, Xiaoyu Wei, Guennadi Kozlov, Yu Chen, Marie Menade

## Abstract

Poly(A) binding protein cytoplasmic 1 (PABPC1) is an essential translational initiation factor. PABPC1 recognizes proteins through conserved PABPC1-interacting motifs 1 and 2 (PAM1 and PAM2). PABPC1-interacting protein-2 (Paip2) interacts with PABPC1 and modulates its activities. Here, we report that the formation of Paip2/PABPC1 complex protects it from proteasome independent degradation. We also show that PAM2 is critical for Paip2/PABPC1 interaction *in vivo*, in agreement with the observation that Paip2 requires PAM2 to interact with PABPC1 on mRNA. Lastly, we propose a role for Paip2 in displacing PABPC1 at the final stage of mRNA deadenylation when the poly(A) tail is partly degraded.

## Introduction

Translation regulation is a critical step in gene expression control. Poly(A) binding protein cytoplasmic 1 (PABPC1) is an essential translation initiation factor and the most abundant cytoplasmic PABP isoform. PABPC1 and eIF4 translation factors prepare mRNA for its recruitment to the translation pre-initiation complex (Sonenberg and Hinnebusch 2009). PABPC1 consists of four RNA-binding domains (RRM1-4) followed by an unstructured linker region and a conserved MLLE domain. The RRM domains mediate the circularization of mRNA through the binding of the 3’ poly(A) tail and eIF4F complex on the mRNA 5’ cap (Imataka, Gradi et al. 1998, Deo, Bonanno et al. 1999, Kahvejian, Svitkin et al. 2005, Safaee, Kozlov et al. 2012). The C-terminal MLLE domain binds a peptide motif, termed PAM2 for <u>PA</u>BPC1-interacting <u>m</u>otif <u>2</u> (Xie, Kozlov et al. 2014).

PAM2-containing proteins, including <u>PA</u>BPC1-<u>i</u>nteracting <u>p</u>rotein 2 (Paip2), can regulate translation and other mRNA related processes. Paip2 was identified based on its interaction with PABPC1 (Khaleghpour, Svitkin et al. 2001). Paip2 inhibits translation *in vitro* through preventing PABPC1 from binding poly(A) RNA and destabilizing the circularized mRNA (Khaleghpour, Kahvejian et al. 2001, Karim, Svitkin et al. 2006). By suppressing PABPC1, Paip2 contributes to control of synaptic plasticity and memory (Khoutorsky, Yanagiya et al. 2013), spermatogenesis (Yanagiya, Delbes et al. 2010), and serves as innate defense to restrict viral protein synthesis to counter the virus-induced increase in PABPC1 (McKinney, Yu et al. 2013).

Paip2 interacts with PABPC1 through two motifs: PAM1 and PAM2. PAM1 binds to the RRM domains of PABPC1, and is characterized by the presence of a large number of negatively charged residues (Khaleghpour, Kahvejian et al. 2001). The binding of PAM1 changes the conformation of PABPC1 and excludes poly(A) RNA binding (Lee, Oh et al. 2014). Interactions between Paip2 PAM2 and PABPC1 MLLE domain are well characterized (Kozlov, De Crescenzo et al. 2004, Kozlov, Menade et al. 2010, Xie, Kozlov et al. 2014). The PAM2 has a highly conserved pattern and a phenylalanine residue is critical for PAM2/MLLE interaction (Kozlov, De Crescenzo et al. 2004). Though the PAM1/RRM and PAM2/MLLE interactions are both observed *in vitro*, it is not clear how the two interaction sites of Paip2/PABPC1 work together in cells.

Here, we report that the PAM2/MLLE interaction is required for Paip2/PABPC1 association *in vivo* and for maintaining Paip2 and PABPC1 stability. Furthermore, we find that Paip2 interacts with PABPC1 on mRNA through PAM2 and could function to displace PABPC1 after shortening of poly(A) tail by deadenylation.

## Results

### Overexpression of PAM2-GFP lowers Paip2 level

To block the Paip2/PABPC1 interaction at the PAM2/MLLE site, we overexpressed in HeLa cells a PAM2 motif of Paip2 (109-125) or a high affinity chimeric superPAM2 (sequence in Materials and Methods, affinity measured in Fig. S1). Cells expressing PAM2-GFP fusion proteins had decreased Paip2 protein levels compared with GFP controls (Fig. 1). This was unexpected, as the two PAM2 motifs also inhibit binding of Paip2 to EDD, a MLLE-containing protein that is involved in ubiquitination and degradation of Paip2 (Yoshida, Yoshida et al. 2006). It was observed but not understood that deletion of PAM2 destabilizes Paip2 due to loss of the interaction with EDD (Yoshida, Yoshida et al. 2006). Our results demonstrate that interaction of Paip2 with the MLLE domain of PABPC1 is important for Paip2 stability.

**Fig. 1.**
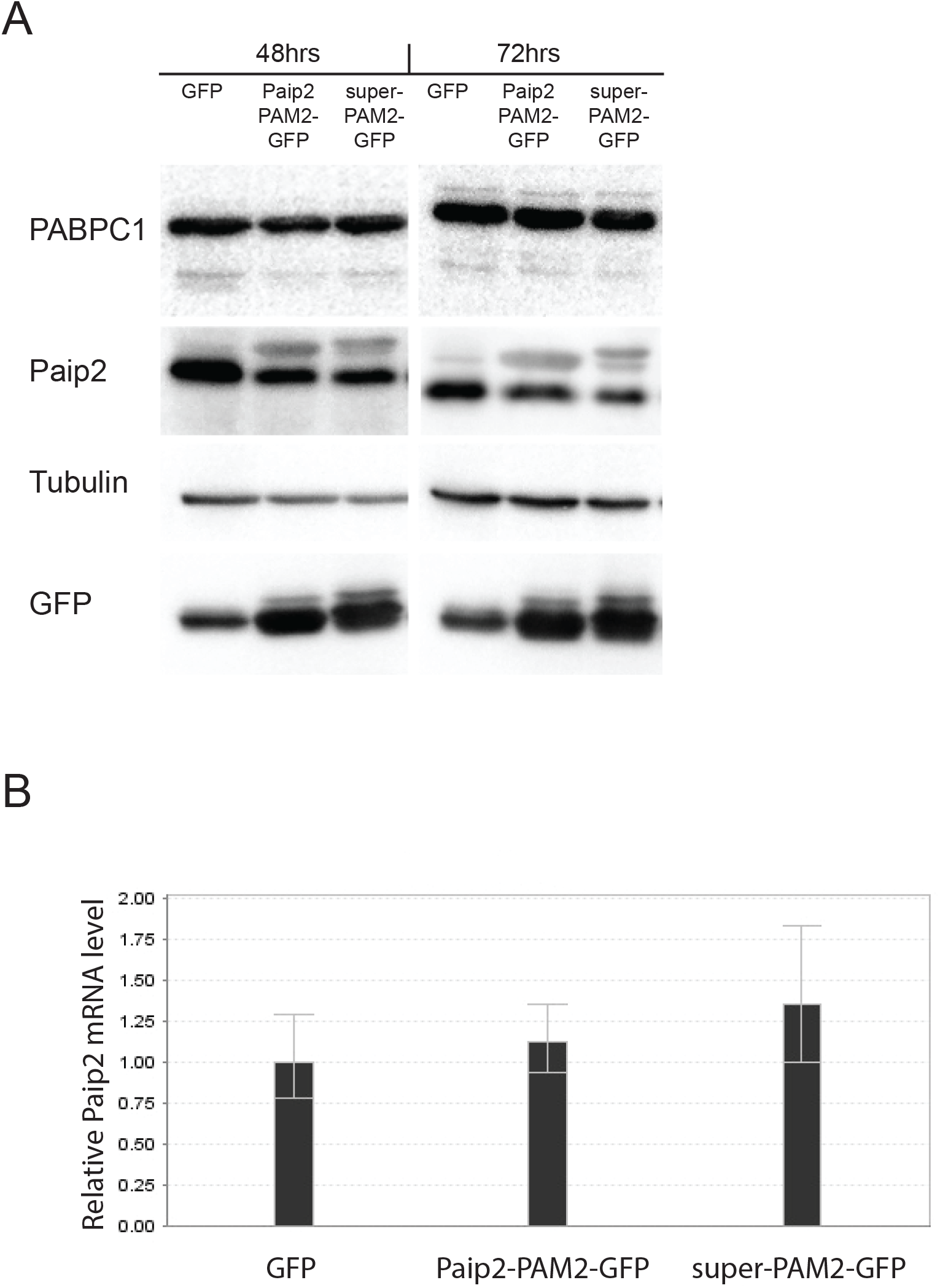
Expression of PAM2-GFP down-regulates Paip2 levels. (A) HeLa cells were transfected with empty vector or constructs expressing GFP tagged Paip2-PAM2 or super-PAM2. Cells were harvested for western blotting after 48 or 72 hours. Reduction of Paip2 levels was observed. (B) Total RNA was extracted with Trizol from cells 72 hours after transfection. qRT-PCR was performed to examine Paip2 mRNA level. mRNA levels were normalized using GAPDH as control.

### Formation of the Paip2/PABPC1 complex protects components from degradation by calpain

Paip2 is subject to calpain-mediated degradation in stimulated neurons independently of the proteasome (Khoutorsky, Yanagiya et al. 2013). To investigate the influence of complex formation on its stability, we treated the complex of full-length, recombinant Paip2 and PABPC1 proteins with calpain I. We found that formation of the complex protected both Paip2 (Fig. 2A) and PABPC1 (Fig. 2B) from degradation by calpain I. Competition from recombinant GST-super-PAM2 added into the reaction mix accelerated Paip2 degradation (Fig. 2). Similarly, PAM2 mutant of Paip2 was degraded faster than wild-type Paip2 (Fig. S2). Interaction with RRM1-4 domains of PABPC1 also protected Paip2 from calpain I digestion. (Fig. S2B). Thus, the protection depended on both PABPC1-binding sites of Paip2. We conclude that the Paip2/PABPC1 interaction prevents protease-mediated degradation of free Paip2 and PABPC1. However, the majority of PABPC1 *in vivo* associates with mRNA (Moretti, Kaiser et al. 2012) and is likely stabilized by this interaction. This explains why PABPC1 protein is not significantly affected by overexpression of GST-PAM2 (Fig. 1).

**Fig. 2.**
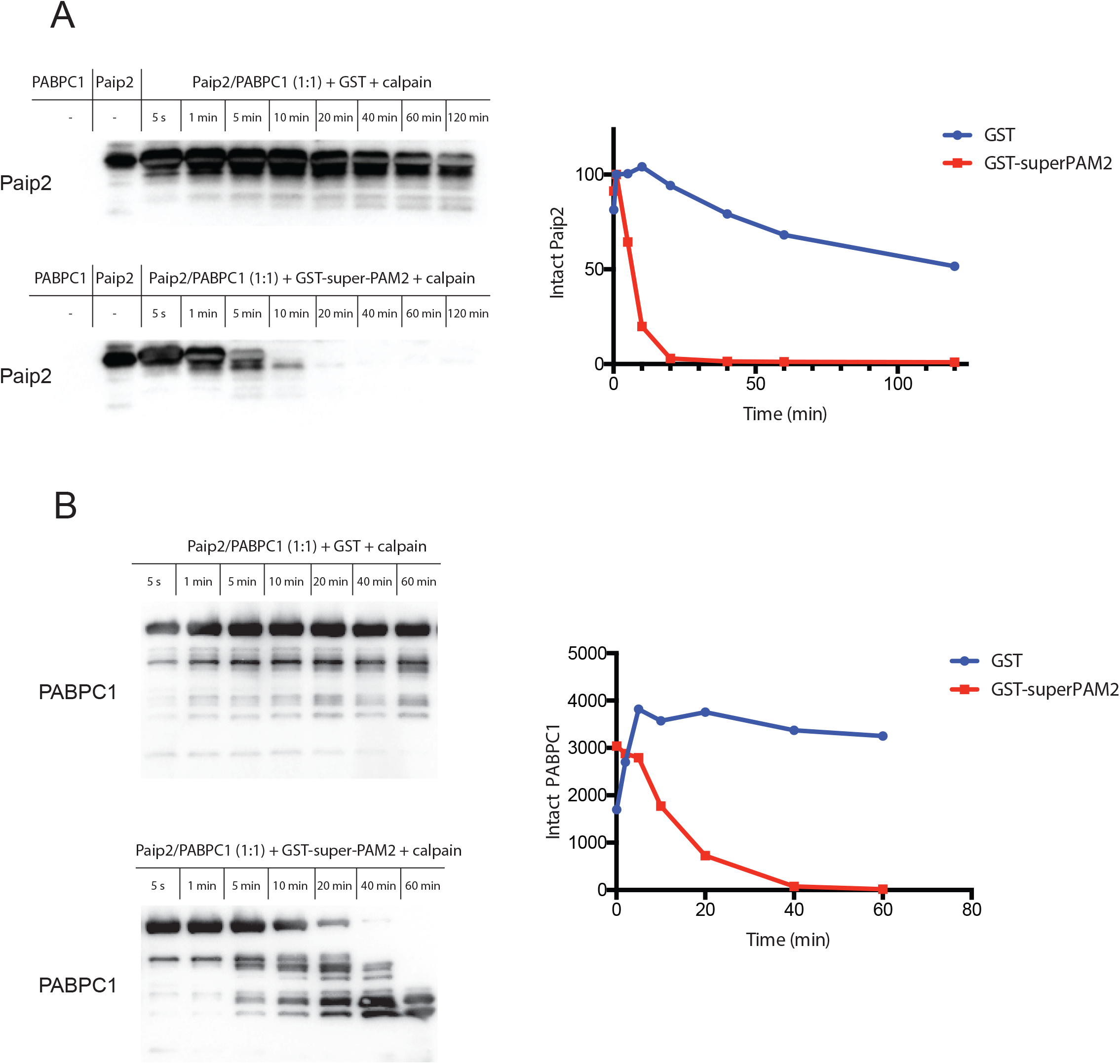
Formation of the Paip2/PABPC1 complex protects components from calpain I degradation. (A) Interaction of Paip2 with PABPC1 through PAM2 binding is important for stability. Paip2 levels decrease following challenge of the Paip2/PABPC1 complex by calpain and GST-super-PAM2. Samples of different time points were collected and the amount of Paip2 analyzed by western blotting and quantified for plotting. (B) PABPC1 levels decrease following challenge of the Paip2/PABPC1 complex by calpain and GST-super-PAM2.

### PAM2/MLLE interaction is critical for Paip2/PABPC1 association in vivo

When the RRM domains of PABPC1 bind mRNA, the MLLE domain is available for binding PAM2-containing proteins, including Paip2 (Xie, Kozlov et al. 2014). Despite the lower affinity of the PAM2/MLLE interaction compared to the PAM1/RRM interaction, we found that the PAM2 motif was required for binding PABPC1 *in vivo*. We transfected cells with GFP-tagged Paip2 or the F118A mutant of Paip2 which does not bind MLLE (Kozlov, De Crescenzo et al. 2004). Overexpressed wild-type or mutant Paip2-GFP was immunoprecipitated by anti-GFP antibody to detect associated endogenous PABPC1. Recombinant GST or GST-super-PAM2 proteins were added to the lysate to test the requirement for the PAM2/MLLE interaction. Although a relatively small population of total PABPC1 was precipitated by GFP-Paip2, it was a representative sample of PABPC1 that bound Paip2. We found that both the PAM2 mutation and competition by GST-superPAM2 significantly decreased the levels of endogenous PABPC1 bound to Paip2 (Fig. 3). This suggests that majority of Paip2 binds PABPC1 in a PAM2-dependent manner *in vivo*. This likely reflects the fact that most PABPC1 is bound to mRNA and, thus, only the PAM2 binding site is available for binding Paip2.

**Fig. 3.**
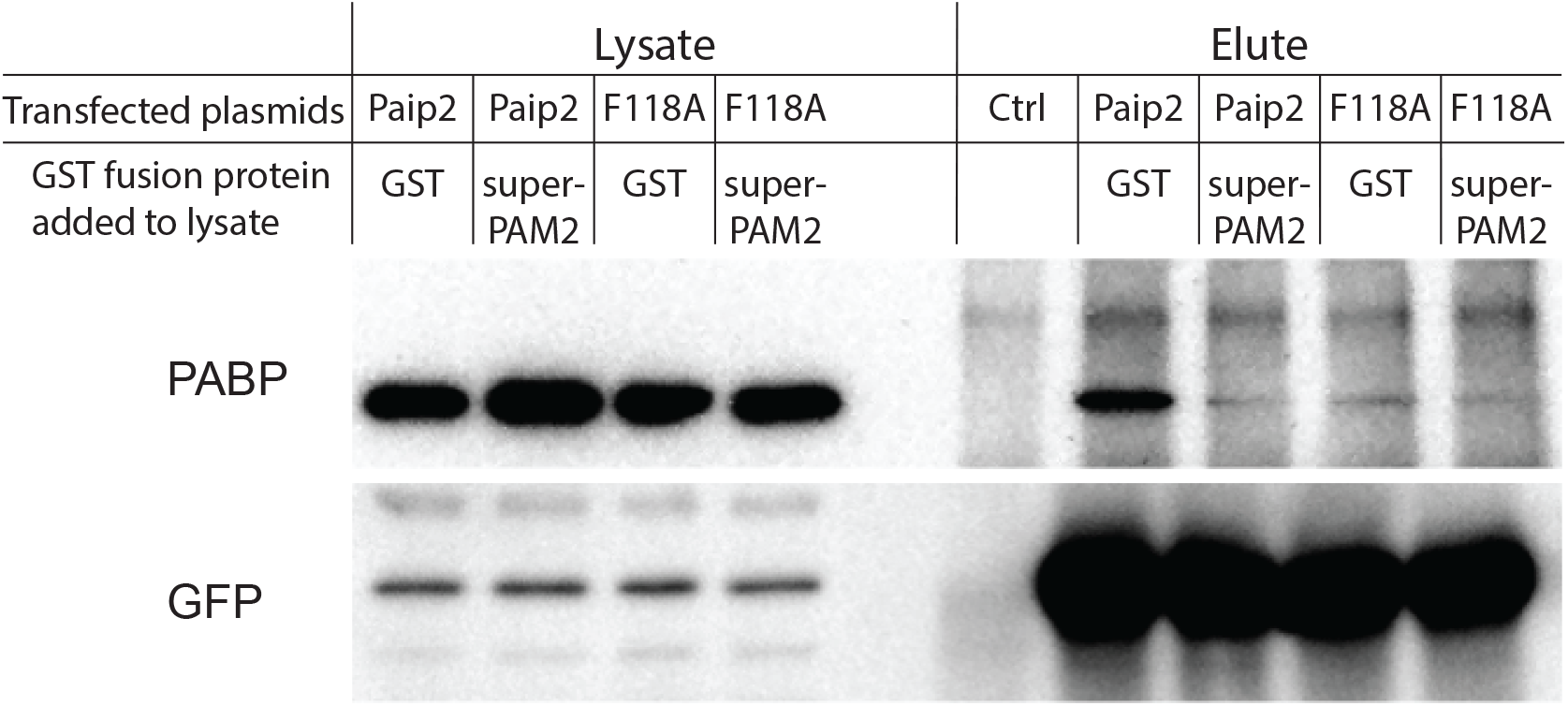
PAM2 is critical for the Paip2/PABPC1 interaction *in vivo*. Paip2-GFP or Paip2(F118A)-GFP were overexpressed in HeLa cells and cell lysates were cleared by centrifugation. GFP antibodies were added to lysate, in presence of GST or GST-super-PAM2 protein. Less PABPC1 was co-immunoprecipitated when the PAM2/MLLE interaction was blocked by mutation of F118 or addition of competing GST-PAM2 protein.

### PABPC1 can interact with Paip2 and mRNA simultaneously

In arsenite-stressed mammalian cells, PABPC1 is enriched in stress granules containing mRNA-protein complexes (Kedersha, Stoecklin et al. 2005). This provides a way to examine whether Paip2 interacts with PABPC1 on mRNA. Paip2 and PABPC1 proteins strongly co-localized at arsenite-induced stress granules (Fig. 4A). To investigate the role of PAM2 in the localization, we transfected cells with Paip2-GFP or Paip2 (F118A)-GFP and induced formation of stress granules. Paip2(F118A) was no longer enriched at stress granules, while wild-type Paip2 (Fig. 4B) and Paip2-GFP were. Thus, Paip2 interacts with PABPC1 on mRNA via interaction with MLLE domain of PABPC1. To visualize the ternary Paip2/PABPC1/RNA complex, we challenged the Paip2/PABPC1 (1:1) complex in vitro with increasing amounts of an RNA oligomer, r(A)_25_. The Paip2/PABPC1/RNA complex displayed a unique migration pattern on native gel. r(A)_25_ readily formed a complex with Paip2/PABPC1 at low molar ratios (Fig. 5A). This implies that RRM domains of PABPC1 prefer binding RNA over Paip2. This agrees with our previous observations that Paip2 needs a functional PAM2 motif to interact with PABPC1 *in vivo* and that PABPC1 is predominantly in complex with mRNA *in vivo*.

**Fig. 4.**
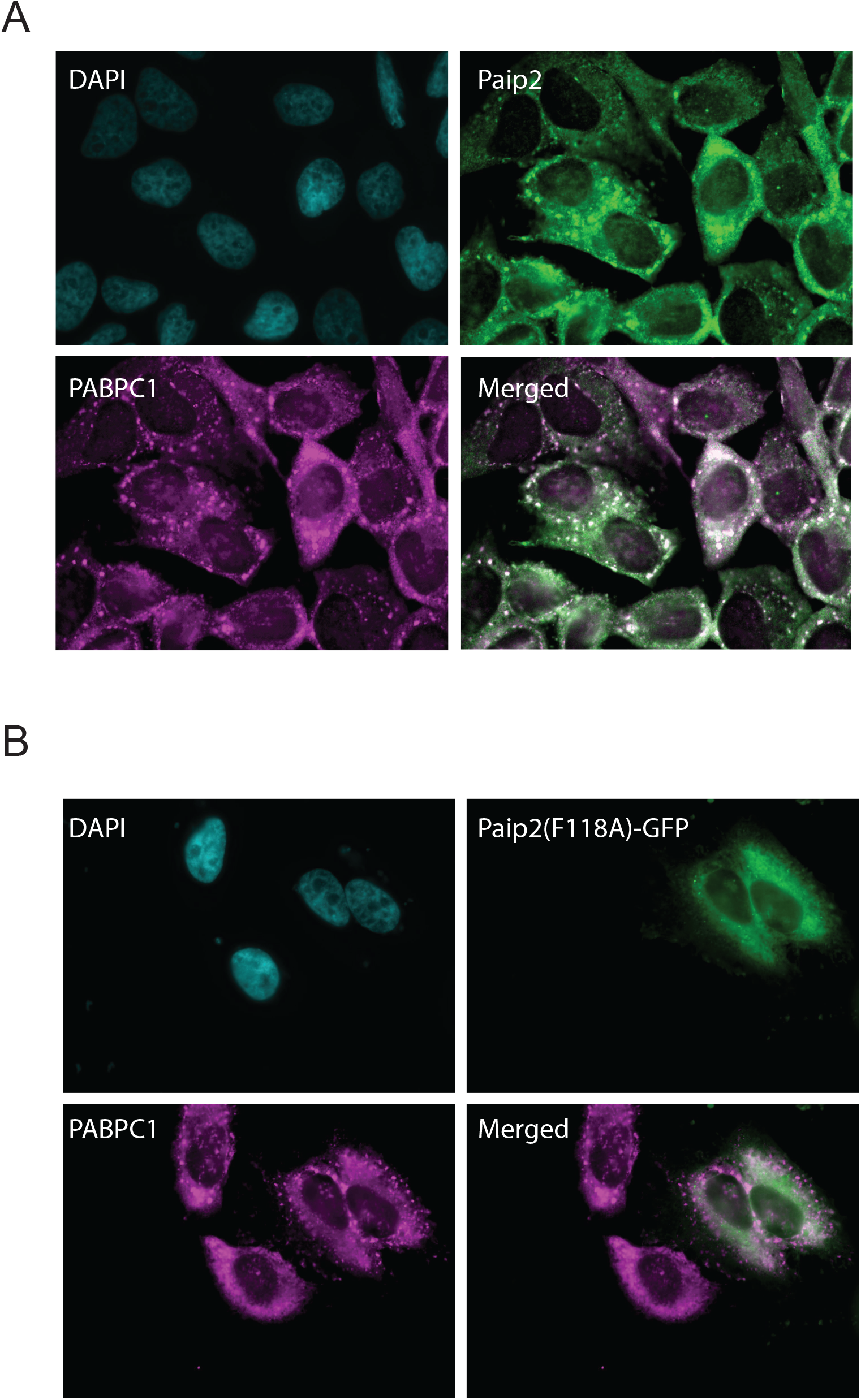
Paip2 localizes to stress granules through interaction with the MLLE domain of PABPC1. (A) HeLa cells were grown on cover slides and fixed after treatment in 0.5 mM sodium arsenite for half an hour. PABPC1 (*magenta*) and Paip2 (*green*) colocalized at stress granules. (B) PAM2 was required for Paip2 localization to stress granules. Cells were transfected with Paip2(F118A)-GFP. The mutant Paip2 did not co-localize with PABPC1.

**Fig. 5.**
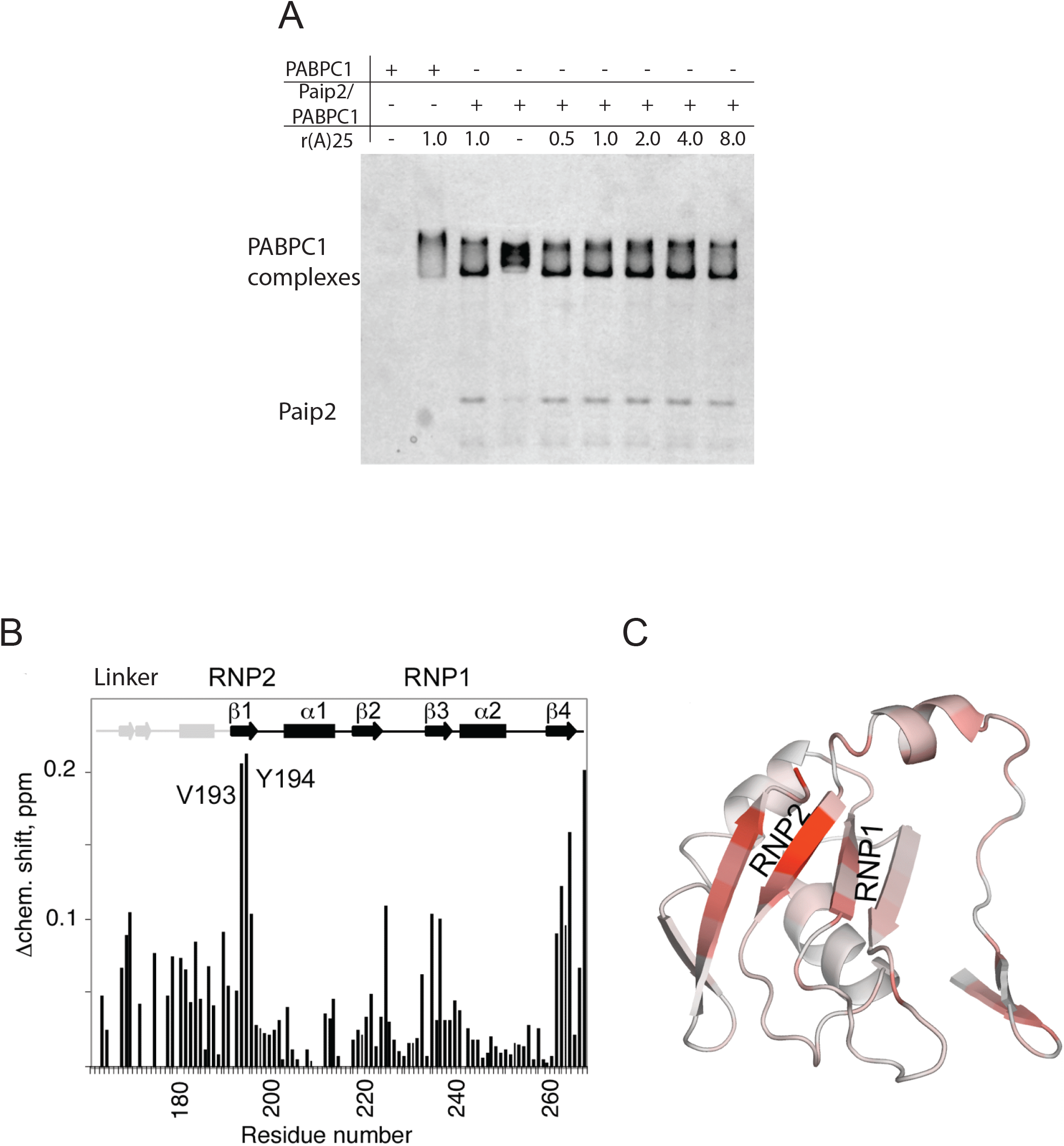
Analysis of Paip2, PABPC1 and poly(A) interplay. (A) Paip2/PABPC1/RNA tertiary complex on native gel. Full-length Paip2/PABPC1 complex were run in native gel. Different molar ratios of r(A)_25_ was added to the protein complex. RNA formed a new complex with Paip2/PABPC1 and gave a new migration pattern. (B) Identification of the Paip2 PAM1 binding site on RRM3 of PABPC1 (166-267). Plot of chemical shift changes in RRM3 upon addition of Paip2 (22-75) identifies regions involved in binding. (C) Color-coded homology model of RRM3 based on the X-ray structure of RRM1-2 (Deo et al, 1999). Residues with large chemical shift changes upon PAM1 addition (panel A) are colored red. The major chemical shift changes are in the RNP2 motif and overlap with the residues predicted to bind RNA.

### PABPC1 uses similar interface on RRM domains to bind Paip2 or poly(A) RNA

Full-length PABPC1 can form a ternary complex with Paip2 and poly(A) RNA as shown in the native gel of Fig. 5A. We then asked how the Paip2 and poly(A) RNA compete with each other for binding to the RRM domains. The N-terminal 75-residue fragment of Paip2A was previously shown to bind to PABPC1 RRM2/3 (Khaleghpour, Kahvejian et al. 2001). NMR experiments with ^15^N-labeled Paip2A (1-75) showed that the N-terminal residues of Paip2A did not participate in the interaction, so a smaller Paip2 (22-75) fragment was cloned and used to study the binding site on PABPC1 (Fig. S3).

Pull-down experiments have shown that, while RRM23 provides full affinity for Paip2, PABPC1 residues R166-Q267 were sufficient to detect binding (Khaleghpour, Kahvejian et al. 2001). This shorter fragment containing RRM3 and the preceding linker sequence gave excellent NMR spectra (Fig. S4). Addition of unlabeled Paip2 (22-75) to ^15^N-labeled PABPC1 (166-267) resulted in specific chemical shift changes for a number of amides indicating binding of the proteins (Fig. S4). The backbone NMR assignments for this PABPC1 fragment alone and in complex with Paip2 (22-75) identified the binding site on the RRM3 domain (Fig. 5B & C). Residues in the β-sheet surface and the RRM2-RRM3 linker showed the largest spectral changes upon Paip2 binding. This same interface is used for recognition of poly(A) RNA (Deo, Bonanno et al. 1999), which explains why the RRM domains of PABPC1 cannot interact with Paip2-PAM1 and poly(A) RNA simultaneously. Thus, when PABPC1 is in complex with mRNA, only the MLLE domain is available for the interaction with Paip2.

### PABPC1 can interact with one or two Paip2 molecules

We have shown that majority of Paip2 interacts with PABPC1 which is binding mRNA through its RRM domains. This suggests that Paip2 does not necessarily bind PABPC1 using both of its binding sites. To characterize the stoichiometry of Paip2/PABPC1 protein complex, we mixed recombinant full-length PABPC1 and Paip2, and analyzed protein complex by size exclusion chromatography-coupled multi-angle light scattering (SEC-MALS). Paip2 and PABPC1 at 1:1 or 2:1 molar ratio both formed mono-dispersed peaks with difference in molecular weight of about 14 kDa, which corresponded to one Paip2 molecule (Fig. 6 A & B). This suggests that PABPC1 can bind either one or two Paip2 proteins. By comparing the root mean square radius (Rg) and the hydrodynamic radius (Rh), we can learn about the compactness of the complex (W. Burchard 1980, S. E. Harding 1992). A higher Rg/Rh ratio indicates an extended conformation. From the Rg/Rh ratio, the Paip2/PABPC1 (2:1) complex was more extended, as PABPC1 was open to bind two Paip2, with one Paip2 at each end (Fig. 6B). Thus Paip2 interacts with PABPC1 at different ratios, depending on the relative abundance of Paip2 and PABPC1 (Fig. 9).

**Fig. 6.**
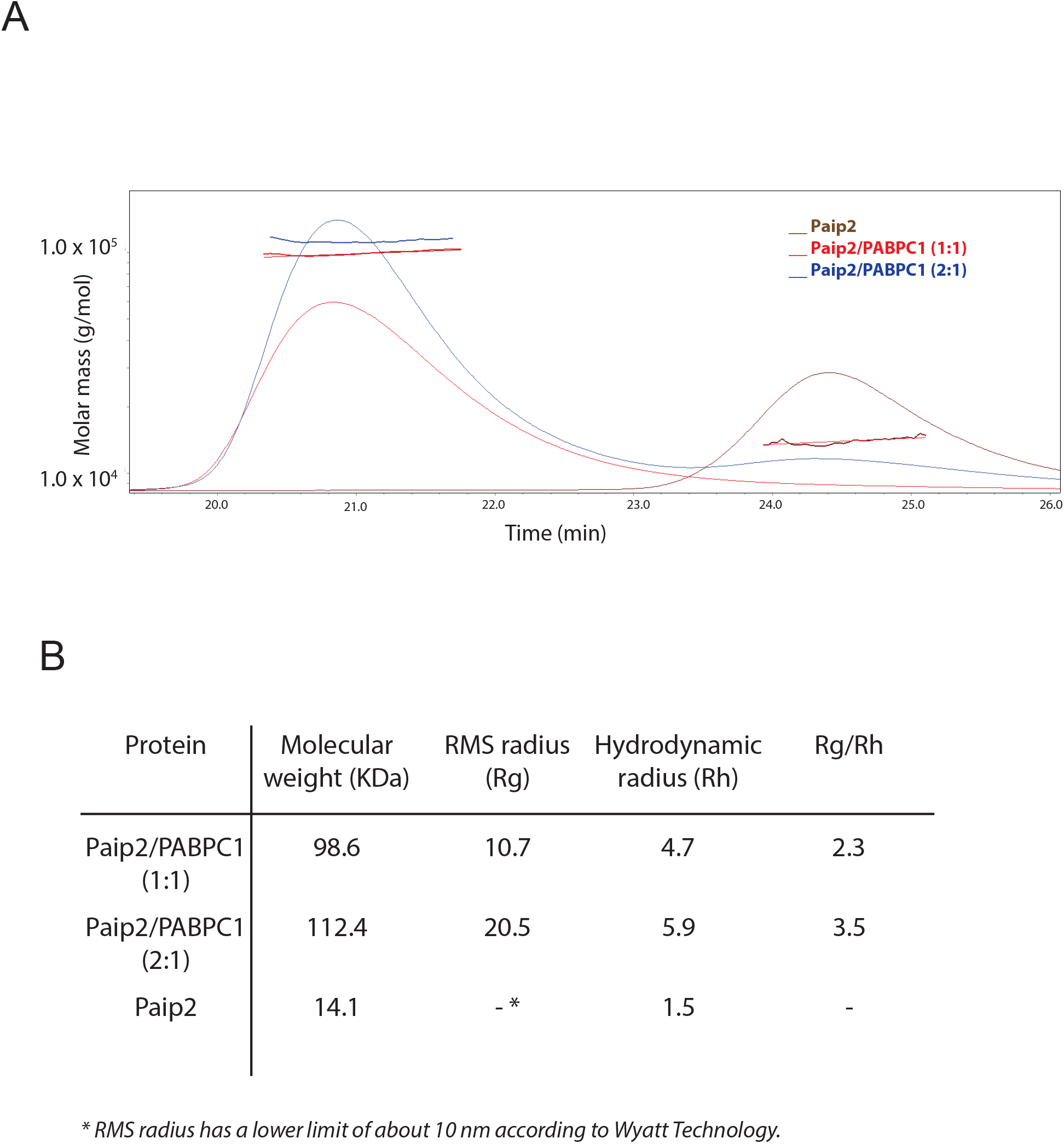
Paip2 and PABPC1 interact with two different ratios. (A) Full-length Paip2/PABPC1 complex was analyzed by SEC-MALS. The molecular weight difference suggested a second Paip2 bound to PABPC1 at ratio 2:1. (B) The calculated molecular weight, root mean square radius and hydrodynamic radius corresponding to Paip2/PABPC1 ratios are listed in the table.

### Paip2 may displace PABPC1 from shortened poly(A) tail of mRNA

The above experiments showed that the RRM domains of PABPC1 prefer to bind to r(A)_25_ over Paip2. To study the interplay of Paip2, PABPC1, and poly(A) RNA, we purified RRM1234 (1-372) of PABPC1 and Paip2-PAM1 (22-75). The two proteins were mixed at 1:1 molar ratio and purified by gel filtration chromatography. The resulting RRM1234/PAM1 complex migrated in a sharp band on non-denaturing PAGE (Fig. 7A). We added increasing r(A)_25_ or r(A)_11_ to the complex. Although the r(A)_25_ or r(A)_11_ has similar affinity for PABPC1 (Sladic, Lagnado et al. 2004), RRM1234/PAM1 was disrupted by r(A)_25_, but remained stable even in the presence of high r(A)_11_ levels (Fig. 7A). This suggested that the length of RNA was critical for competing with Paip2. To further understand the competition, we assembled a complex of RRM23 (98-293) and Paip2 (22-75), and tested whether different RNAs could disrupt the protein complex (Fig. 7B & C). We found r(A)_16_ disrupted the RRM23/Paip2 complex, while r(A)_11_, (AUUU)_3_, and (AUUU)_4_ did not. Although PABPC1 binds all these RNA oligos *in vitro*, the length and sequence requirements of RNA in disassembly of RRM23/Paip2 indicate that longer poly(A) RNAs induce a unique conformation of PABPC1 that disfavors Paip2.

**Fig. 7.**
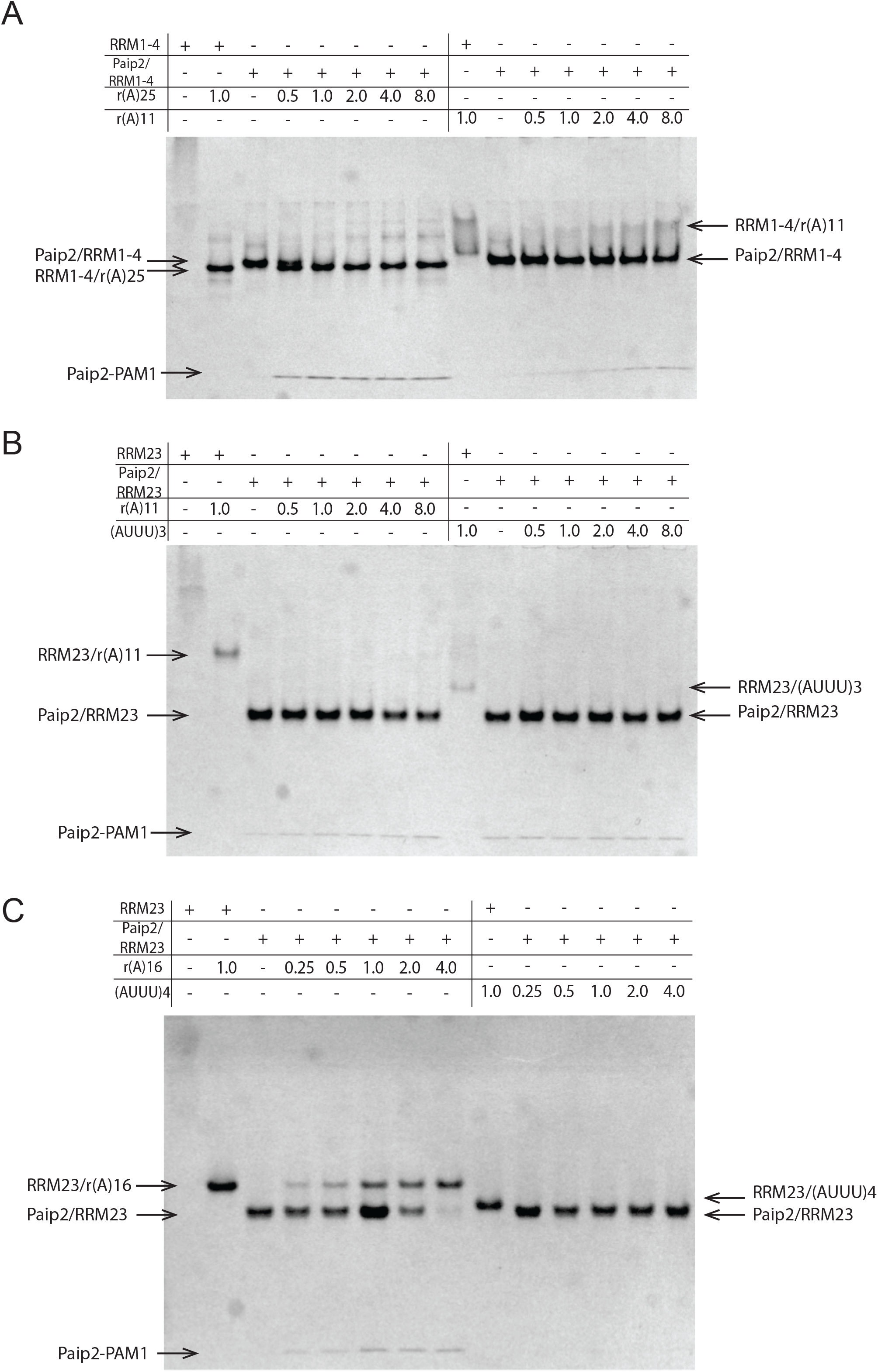
PABPC1 prefers binding to Paip2 over short poly(A) or (AUUU) repeats. (A) r(A)_25_ disrupts RRM1234/Paip2 complex. 60 pmol of PABPC1 (1-372) (or RRM1234)/Paip2 (22-75) complex were challenged with increasing amounts of r(A)_25_ or r(A)_11_. The molar ratios of RNA compared to PABPC1/Paip2 complex are indicated above the gel. (B) r(A)_11_ or (AUUU)_3_ could not disrupt RRM23/Paip2 complex. (C) r(A)_16_ disrupted RRM23/Paip2 complex, while (AUUU)_4_ did not.

### Depletion of Paip2 reduced microRNA-mediated gene silencing

The change of PABPC1’s preference towards Paip2 with the shortening of RNA can be of great significance in the displacement of PABPC1 during mRNA deadenylation (Fig. 9). We examined the effects of Paip2 depletion in a Renilla and firefly dual luciferase reporter assay (Pillai, Bhattacharyya et al. 2005). Presence of let-7 microRNA binding sites silenced the reporter by about 10 times, and the silencing was significantly attenuated after Paip2 knockdown (Fig. 8A & B). Consistent with our results, it was reported by two groups that Paip2 knockdown diminished microRNA-mediated silencing activity (Walters, Bradrick et al. 2010, Yoshikawa, Wu et al. 2015). Given the fact that very high concentration of Paip2 is required to disrupt the PABPC1/poly(A) RNA interaction *in vitro* (Khaleghpour, Svitkin et al. 2001), we hypothesize that Paip2 does not actively compete with poly(A) for PABPC1, but instead enhances deadenylation at a later stage by promoting dissociation of PABPC1 from the shortened mRNA. Knockdown of Paip2 reduced microRNA-mediated silencing in luciferase reporter assay, likely through inhibition of late steps in the deadenylation process.

**Fig. 8.**
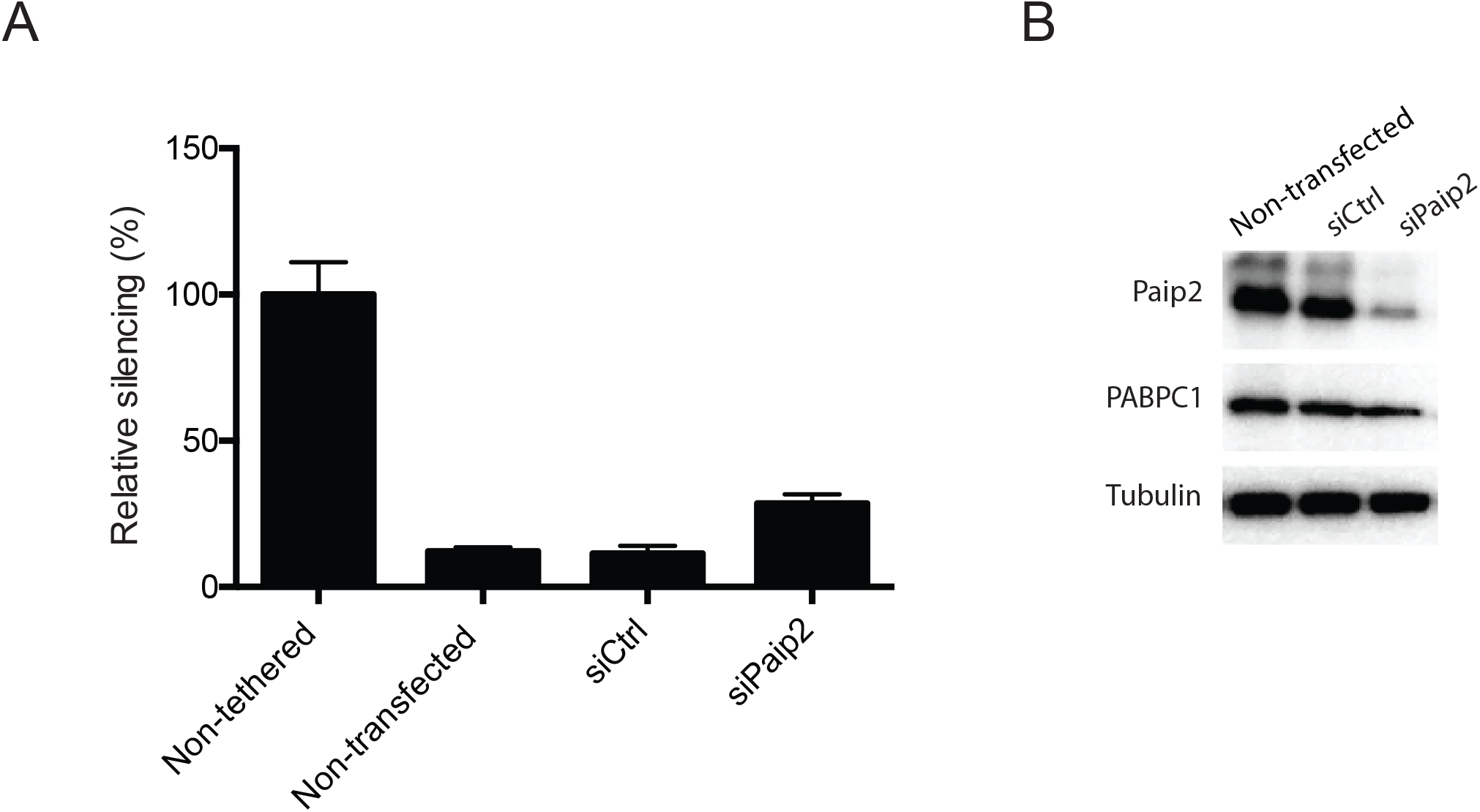
Knockdown of Paip2 reduced microRNA-mediated gene silencing. (A) HEK 293T cells were co-transfected with siRNA control or siPaip2 and firefly luciferase (FL) and Renilla luciferase (RL) plasmids. Luciferase activities from plasmids with three let-7 binding sites or mutated binding sites were normalized to firefly luciferase activities. The reduction of silencing activity by siPaip2 was significant at confidence level of 0.05, by two-tail t-test for two samples with unequal variances. (B) Paip2, PABPC1 and tubulin levels were probed by western blotting in cells after the same treatment as in (A).

## Discussion

In this study, we illustrate a novel role of Paip2 in mRNA metabolism. Our results demonstrate that Paip2 interacts with PABPC1 on mRNA and can exclusively bind PABPC1 when the poly(A) tail is shortened. This supports the model that Paip2 enhances mRNA deadenylation through facilitating PABPC1 dissociation.

PABPC1 is required for deadenylation of mRNA by Pan2/Pan3 deadenylases to a short length of about 10-25 nucleotides (Lowell et al., 1992; Zheng et al., 2008). At that point, the mRNA is stabilized and protected by PABPC1 from further deadenylation by Ccr4/Not deadenylases (Tucker et al., 2002). The last PABPC1 molecule needs to be displaced at this stage to finish deadenylation. However, the displacement of PABPC1 from mRNA is poorly understood. Paip2 has been known to act as a competitor of RNA for PABPC1, but very high concentrations of Paip2 are needed to disrupt the PABP/r(A)_25_ complex (Khaleghpour, Svitkin et al. 2001). Here, we show that Paip2 associates with PABPC1 on mRNA and out-competes shortened poly(A) RNA for binding to PABPC1, which renders Paip2 a good candidate as a mediator of the final displacement (Fig. 9). Nonetheless, other mechanisms for dissociate PABPC1 off mRNA remain possible. In microRNA-mediated gene silencing, PABPC1 facilitates recruitment of GW182-containing silencing complex and later appears to be displaced prior to deadenylation (Moretti, Kaiser et al. 2012, Zekri, Kuzuoglu-Ozturk et al. 2013). Multiple mechanisms may come into play for different scenarios in cells.

**Fig. 9.**
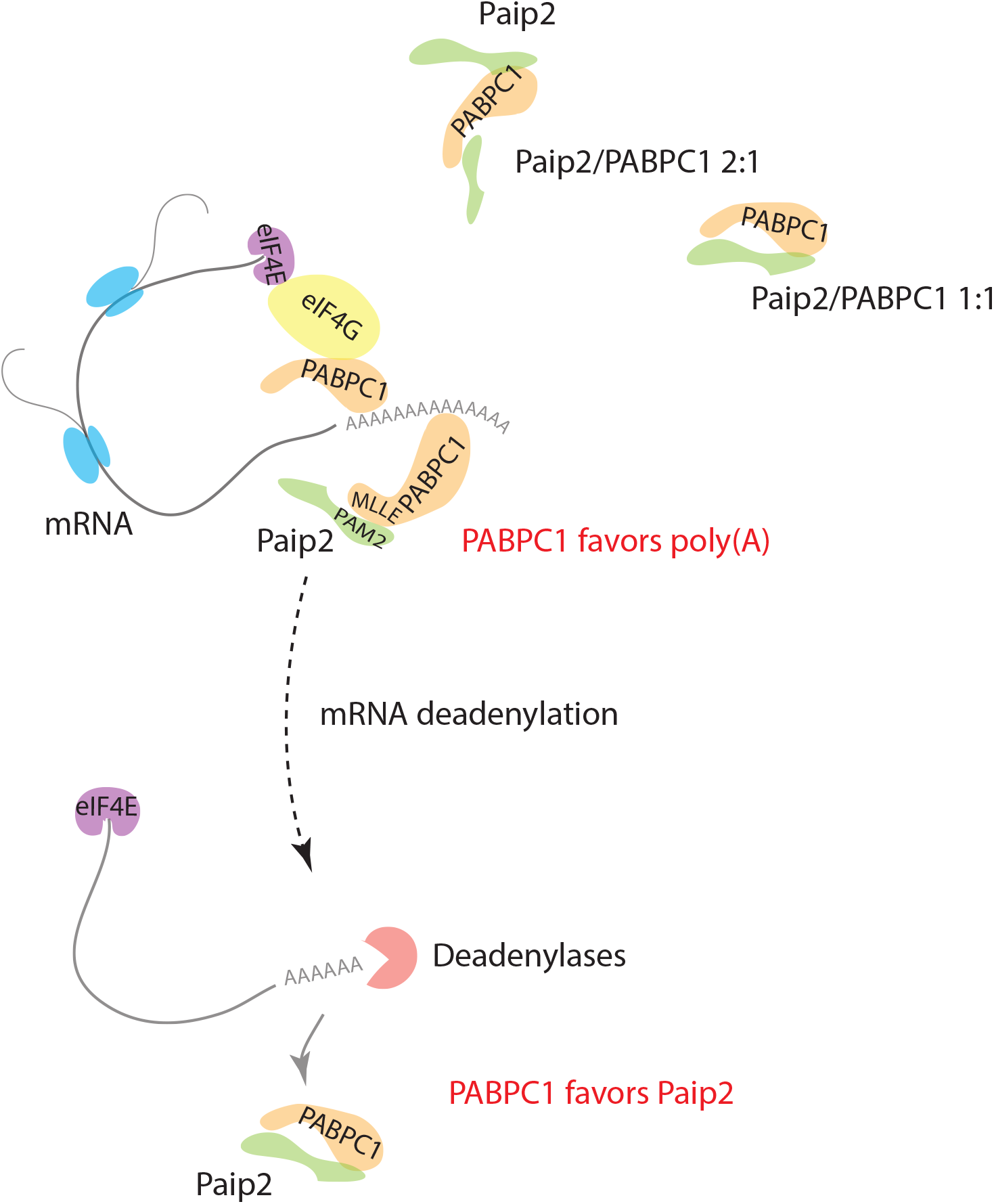
Model of the Paip2 and PABPC1 interaction in cells. Paip2 associates with PABPC1 on mRNA. In the cytoplasm, Paip2 may interaction with PABPC1 with stoichiometries of 1:1 or 2:1. When mRNA poly(A) tail is shortened by deadenylation, Paip2 may actively displace PABPC1 from the mRNA and protect PABPC1 by formation of a protein complex.

PABPC1 has a flexible linker in its middle region, which is targeted by a number of proteases (Smith and Gray 2010). Immediate binding of PABPC1 to Paip2 after dissociation from mRNA offers protection to mRNA-free PABPC1 in cytoplasm. Indeed, addition of Paip2 slows down PABPC1 cleavage by 3C and 2A (Rivera and Lloyd 2008). The interaction between Paip2 and PABPC1 on mRNA makes the immediate protection possible.

Less than 10% of PABPC1 can be depleted by GST-Paip2 without micrococcal nuclease treatment (Moretti, Kaiser et al. 2012), which suggests that over 90% of PABPC1 in cells is bound to mRNA. According to our data, the levels of PABPC1 are 2 to 4 times higher than Paip2 in HeLa, HEK293 or MEF cells (Fig. S5). Thus, there is an excess of Paip2 in cytoplasm relative to free PABPC1 not associated with mRNA. This is supported by the co-localization of Paip2 with PABPC1 at stress granules (Fig. 4).

Paip2 is subjected to extensive proteasome-dependent (Yoshida, Yoshida et al. 2006) and proteasome-independent (Khoutorsky, Yanagiya et al. 2013) degradation. It is conceivable that regulation of Paip2 levels is used by host to restrict viral protein synthesis (McKinney, Yu et al. 2013). An optimal level of Paip2 is important for spermatogenesis (Yanagiya, Delbes et al. 2010) and microRNA-mediated gene silencing (Walters, Bradrick et al. 2010, Yoshikawa, Wu et al. 2015).

In conclusion, we have shown that Paip2 uses its dual interaction sites to interact with PABPC1 in cells. Paip2 protects free PABPC1 and may actively function through PABPC1 on mRNA. More studies are needed to elucidate the upstream and downstream pathways of this Paip2 and PABPC1 interplay.

## Materials and methods

### Plasmids and RNA oligos

pET-28B-PABPC1 (NM_002568.3) plasmid was a gift from Drs. Svitkin and Sonenberg at McGill University. Firefly luciferase and renilla luciferase reporter plasmids were kindly provided by Dr. Filipowicz at the Friedrich Miescher Institute for Biomedical Research. PABPC1 (1-372), Paip2a (NM_001033112.2), Paip2a-PAM2 (109-125), Paip2a-PAM1 (22-75) were cloned into pGEX-6p1 for protein expression in *E. coli*. Paip2a-PAM2 (109-125) and super-PAM2 (SNLNPNAPEFHPGVPWKGLQNI) were cloned into pCDNA3.1–EGFP. Paip2a was subcloned into pCDNA3.1-EGFP and mutated to express Paip2aPhe118Ala-GFP, with QuikChange site-directed mutagenesis kit (Agilent). siPaip2 (5’-GAGUACAUGUGGAUGGAAAUU-3’), RNA oligo r(A)_25_ and r(A)_11_ were synthesized, desalted and gel purified at Dharmacon. Control siRNA was purchased from Qiagen (SI03650318).

### Protein purification

BL21 (DE3) *E. coli* was transformed with related plasmids and induced with 1 mM IPTG at an OD of 0.7. Bacteria were grown for 3 hours at 30 degrees before being harvested. Pelleted cells were resuspended in 1x PBS buffer (pH 7.4) with 5 mM BME and proteinase inhibitor cocktail (Roche). For purification of PABPC1-His, 10 mM imidazole, 10 μg/ml RNase A and 10 μg/ml DNase I were added to lysate. PABPC1-His was bound to Ni-NTA resin (Qiagen), washed by 1x PBS (10 mM imidazole added) and eluted with 200 mM imidazole. PABPC1 (1-372) was purified using GST resin (GE) and cleaved on the column by 3C PreScission protease. Eluted PABPC1-His or PABPC1 (1-372) was run through anion-exchange and cation-exchange columns (Biosuite Q and Biosuite SP, 13 μm, 21.5 x 150 mm column, Waters), before it was finally applied to Superdex 200 Hiload 16/60 size-exclusion column (GE). The buffer for gel filtration was 20 mM HEPES pH 7.0, 150 mM NaCl, 2 mM DTT. Paip2 was purified using Superdex 75 after affinity purification by GST tag and cleavage on the column. The PABPC1 (1-372)/PAM1 complex was obtained by incubation of purified RRM1234 and 1.5 fold molar excess of GST-Paip2 (22-75) in 1x PBS supplemented with 3C protease and 2 mM DTT. The protein complex was further purified using Superdex 75 Hiload 16/60 column (GE).

### Cell culture and transfection

HeLa cells were cultured in DMEM supplemented with antibiotics and 10% fetal bovine serum. 10^5^ cells per well were plated in 24-well plate the day before transfection. 0.8 μg DNA plasmid was mixed with 2 μl Lipofectamine 2000 in Opti-MEM and then added to cells. After 24 hours, cells were trypsin digested and split onto cover slides. Cells were treated with 0.5 mM sodium arsenite for 0.5 hour and fixed on the second day.

### Immunofluorescence and confocal microscopy

Cells were fixed with 4% PFA in 1x PBS, and penetrated by cold methanol (−20°C) or 0.1% Triton X-100 in 1x PBS for 10 minutes. Cells were blocked with 5% normal goat serum (Millipore S26) in PBS for 1 hour. Then cells were incubated in 1x PBS, supplemented with anti-PABPC1 (Abcam ab21060; Santa Cruz sc32318)(1:200) or anti-Paip2 (Sigma-Aldrich P10087). Cells were washed in PBS 3 times, before being incubated with corresponding second antibodies conjugated with Dylight550 (Bethyl A120-101D3) or Rhodamine (Millipore 12-509, 12-510) at 1:200 – 1:500 dilutions. DAPI (Roche) was added to washing buffer at 0.5 μg/ml to treat cells for 10 minutes. Cover slides were finally mounted in ProLong Gold antifade reagent (Life Technology P36930). Images were collected on Zeiss LSM 310 confocal microscope in the McGill University Life Sciences Complex Advanced BioImaging Facility (ABIF).

### Quantitative RT-PCR

Total RNA was extracted from cells with Trizol (Life Technology). cDNA libraries were prepared using SuperScript First-Strand Synthesis System for RT-PCR (Life Technology). Validated Taqman assays were purchased for quantification of GAPDH (Applied Biosystems Hs 99999905) and Paip2 (Applied Biosystems Hs00212868). qRT-PCR reactions were run in Stepone Plus PCR system (Applied Biosystems).

### Immunoprecipitation

Cells were lysed in 20 mM HEPES (pH 7.4), 150 mM NaCl, 0.5% NP-40, 2 mM DTT, 2 mM MgCl_2_, 1 mM CaCl_2_ and protease inhibitor tablet (Roche), and cleared by centrifugation. 2 μg antibodies were added per 1 mg cleared lysate for 2 hours. Dynabeads conjugated with protein A or protein G (Life Technology) were washed and added to lysate for 0.5 hour. Dynabeads were then washed with 1x PBS and boiled in SDS loading buffer for further analysis.

### Western blotting

Protein samples were heated at 95°C and separated by SDS-PAGE. Proteins were then transferred to PVDF membrane (Millipore) in Tris/Glycine buffer with 20% methanol in cold room. PVDF membrane was blocked in 1x TBST (pH 7.5), containing 0.05% Tween-20 and 5% skim milk powder or bovine serum albumin. The membrane was then incubated with primary antibodies, including anti-PABPC1 (Abcam ab21060 1:1000), anti-Paip2 (Sigma-Aldrich P10087 1:2000), anti-tubulin (Sigma-Aldrich T9028 1:5000) and anti-GFP (Clontech 632381 1:2000). Membrane was then washed three times in 1x TBST and incubated with goat-anti-rabbit (Jackson ImmunoResearch 111-035-046 1:5000) or goat-anti-mouse (Jackson ImmunoResearch 115-035-071 1:5000) for 0.5 hour. After wash, membrane was developed with Amersham ECL prime kit (GE healthcare RPN2236) and imaged with Alpha Innotech imaging system.

### NMR spectroscopy

NMR samples were prepared in 90% NMR buffer (10 mM HEPES pH 7.0 and 50 mM NaCl) and 10% D_2_O. NMR resonance assignments of the RRM3 domain were carried out using standard heteronuclear 3D-experiments HNCACB and CBCA(CO)NH on ^13^C,^15^N-labeled protein. For NMR titrations, unlabeled Paip2 (22-75) was added stepwise to 0.2 mM ^15^N-labeled RRM3 to a final molar ratio of approximately 3 to 1. Chemical shift changes were calculated according to the formula 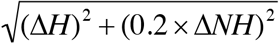. All NMR experiments were performed at 301 K on a Bruker 600 MHz spectrometer. NMR spectra were processed using NMRPipe(Delaglio, Grzesiek et al. 1995) and analyzed with XEASY(Bartels, Xia et al. 1995).

### Size Exclusion Chromatography-coupled Multi-Angle Light Scattering

200 μg of recombinant PABPC1 protein was mixed with Paip2 at indicated molar ratios. Protein mixture was injected into Superdex 200 10/300 GL column and analyzed by a MiniDAWN TREOS light-scattering detector (Wyatt Technology Corporation) and a Optilab rEX (Wyatt) refractive index detector. All analysis was done in 20 mM HEPES pH 7.0, 150 mM NaCl, 2 mM DTT. Data processing was performed using Astra software (Wyatt).

### Fluorescence polarization assay

Peptides SNLNPNAPEFHPGVPWKGLQNI-FITC (super-PAM2) and SNLNPNAPEAHPGVPWKGLQNI-FITC (super-PAM2-F10A) were synthesized at EZBiolab. 5 nM of fluorophore-labeled peptide was mixed with serial dilutions of GST-PABPC1-MLLE or GST-EDD-MLLE. Fluorescence polarization signals of each well were recorded on a SpectraMax M5e Microplate Reader at excitation wavelength of 495 nM and emission wavelength of 519 nM. Fluorescence polarization signals as milli-polarization units (mP) were plotted against a log scale of protein concentration (μM).

### Calpain digestion of proteins

40 μg Paip2 was mixed with equal molar amount of PABPC1 and then incubated with Calpain I. Samples were collected at indicated time points. GST or GST-super-PAM2 was added as a control or to compete with Paip2 for interaction with PABPC1.

Degradation of Paip2 was examined by western blotting using anti-Paip2 or anti-PABPC1 antibodies. Bands were quantified in Alphaview software (AlphaInnotech), and plotted against digestion time.

### Dual luciferase reporter assay for microRNA-mediated silencing activity

Renilla and firefly luciferase reporters were described before(Pillai, Bhattacharyya et al. 2005). For overexpression assays, human HeLa cells were seeded in 24-well plates and transfected using Lipofectamine 2000 (Life Technology). The transfection mixtures contained 10 ng of R-Luc-3xlet-7 reporter plasmid (RL-3xlet-7) or the corresponding reporter carrying mutations in the let-7-binding sites (RLuc-Mut), 100 ng of the pEGFP-N3-F-Luc transfection control and 60 pmol of siRNA control or siPaip2. R-Luc and F-Luc activities were measured 48 h after transfection using the Dual-Luciferase Reporter Assay System (Promega). Renilla luciferase activity was normalized to firefly luciferase.

## Acknowledgements

We are grateful for suggestions and help from colleagues during the project.

## Author contributions

J. X. and G. K. designed the experiments. J. X., W. X., G. K., and Y. C. carried out the experiments. J. X., G. K., and K. G. wrote the manuscript.

---

<u>Fig. S1</u>. Affinities of <u>super-PAM2 to MLLE domain of PABPC1 and EDD</u> measured by fluorescence polarization. Peptides were <u>labeled with FITC</u> and fluorescence measured in the presence of the MLLE domains of PABPC1 or EDD.

<u>Fig. S2. Paip2 degradation by calpain I</u>. (A) Paip2-PAM2 mutant (F118A) was degraded faster compared to wild-type Paip2 in presence of PABPC1. (B) Paip2 or Paip2F118A were rapidly degraded with equal rates by calpain I.

<u>Fig. S3. Mapping of PAM1 region of Paip2 to residues 22-75</u>. 15N-HSQC spectra of Paip2 (1-75) before and after addition of RRM23 are shown side by side. Upon addition of RRM23 fragment of PABPC1 to ^15^N-labeled Paip2 (1-75), signals of the residues involved in PABPC1 interactions undergo broadening, which results in weak or disappearing signals. In contrast, residues that do not bind to PABPC1 produce sharp signals and can be identified using 3D ^15^N-TOCSY experiment. In particular, all serine and glycine residues (labeled as S and G, respectively) reside in the N-terminal 22-residue fragment of Paip2 (1-75) and do not interact with PABPC1 RRM23, while side chains (Ws.ch.) of all three tryptophans (W36, W52, W73) are in the bound region of Paip2.

<u>Fig. S4. Overlay of the ^1^H-^15^N correlation NMR spectra of the ^15^N-labeled PABPC1 RRM3 domain (residues 166-277) alone (in black) and in the presence of equimolar amount of unlabeled Paip2 (22-75) (in green)</u>. NMR titration resulted in specific chemical shift changes indicating binding between the proteins.

<u>Fig. S5. Quantification of amounts of PABPC1 and Paip2 in cells</u>. (A) Dilutions of recombinant PABPC1 or Paip2 protein were analyzed in SDS-PAGE together with cytoplasmic extract (CE) or nuclear extract (NE) of HeLa, MEF, and HEK 293T cells. Proteins were detected by western blotting using anti-PABPC1 or anti-Paip2 antibodies. (B) Signals of purified PABPC1 or Paip2 were plotted against loaded concentrations to make standard curves. Relative amounts of PABPC1 and Paip2 were determined by fitting signal intensities into the curve. All the cytoplasmic extract signals fell into the linear ranges of standard curves. PABPC1/Paip2 ratios were calculated as in the table. (C) Coomassie blue staining of purified PABPC1 and Paip2 proteins. PABPC1 and Paip2 were run in SDS-PAGE. Paip2 shows retarded migration in SDS-PAGE, due to the high content of acidic amino acids.

