## Supplemental figures for "Paip2 associates with PABPC1 on mRNA, and may facilitate PABPC1 dissociation from mRNA upon deadenylation"

Fig. S1. Affinities of super-PAM2 to MLLE domain of PABPC1 and EDD measured by fluorescence polarization. Peptides were labeled with FITC and fluorescence measured in the presence of the MLLE domains of PABPC1 or EDD.

Fig. S2. Paip2 degradation by calpain I. (A) Paip2-PAM2 mutant (F118A) was degraded faster compared to wild-type Paip2 in presence of PABPC1. (B) Paip2 or Paip2F118A were rapidly degraded with equal rates by calpain I.

Fig. S3. Mapping of PAM1 region of Paip2 to residues 22-75. 15N-HSQC spectra of Paip2 (1-75) before and after addition of RRM23 are shown side by side. Upon addition of RRM23 fragment of PABPC1 to $^{15}$N-labeled Paip2 (1-75), signals of the residues involved in PABPC1 interactions undergo broadening, which results in weak or disappearing signals. In contrast, residues that do not bind to PABPC1 produce sharp signals and can be identified using 3D $^{15}$N-TOCSY experiment. In particular, all serine and glycine residues (labeled as S and G, respectively) reside in the N-terminal 22-residue fragment of Paip2 (1-75) and do not interact with PABPC1 RRM23, while side chains (Ws.ch.) of all three tryptophans (W36, W52, W73) are in the bound region of Paip2.

Fig. S4. Overlay of the $^{1}$H-$^{15}$N correlation NMR spectra of the $^{15}$N-labeled PABPC1 RRM3 domain (residues 166-277) alone (in black) and in the presence of equimolar amount of unlabeled Paip2 (22-75) (in green). NMR titration resulted in specific chemical shift changes indicating binding between the proteins.

Fig. S5. Quantification of amounts of PABPC1 and Paip2 in cells. (A) Dilutions of recombinant PABPC1 or Paip2 protein were analyzed in SDS-PAGE together with cytoplasmic extract (CE) or nuclear extract (NE) of HeLa, MEF, and HEK 293T cells. Proteins were detected by western blotting using anti-PABPC1 or anti-Paip2 antibodies. (B) Signals of purified PABPC1 or Paip2 were plotted against loaded concentrations to make standard curves. Relative amounts of PABPC1 and Paip2 were determined by fitting signal intensities into the curve. All the cytoplasmic extract signals fell into the linear ranges of standard curves. PABPC1/Paip2 ratios were calculated as in the table.

(C) Coomassie blue staining of purified PABPC1 and Paip2 proteins. PABPC1 and Paip2 were run in SDS-PAGE. Paip2 shows retarded migration in SDS-PAGE, due to the high content of acidic amino acids.

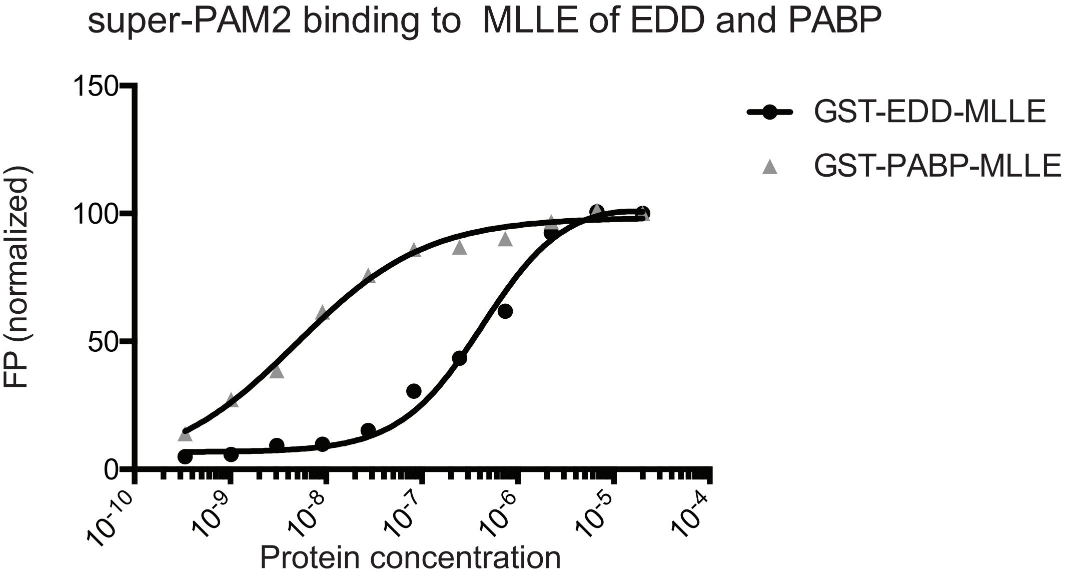


Affinities of PAM2s to MLLE domains

| | PABPC1-MLLE | EDD-MLLE |
|---|---|---|
| PAM2 (Paip2) | 0.12 μM | 6 μM |
| super-PAM2 | 0.03 μM | 0.2 μM |

Fig. S2

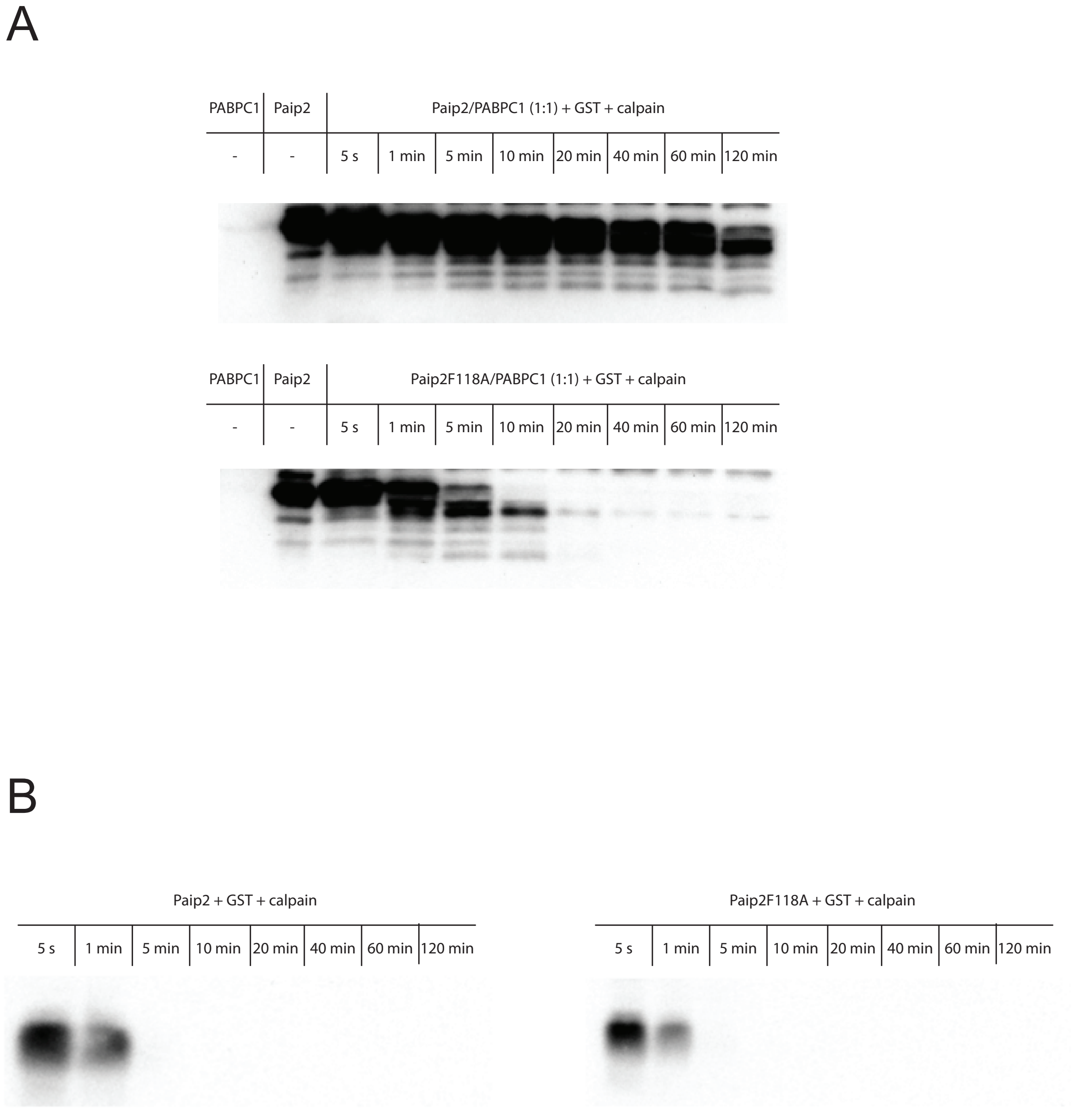
A
PABPC1
Paip2
Paip2/PABPC1 (1:1) + GST + calpain
-
-
5 s
1 min
5 min
10 min
20 min
40 min
60 min
120 min
PABPC1
Paip2
Paip2F118A/PABPC1 (1:1) + GST + calpain
-
-
5 s
1 min
5 min
10 min
20 min
40 min
60 min
120 min
B
Paip2 + GST + calpain
5 s
1 min
5 min
10 min
20 min
40 min
60 min
120 min
Paip2F118A + GST + calpain
5 s
1 min
5 min
10 min
20 min
40 min
60 min
120 min

Fig. S3

HSQC of Paip2 (1-75)

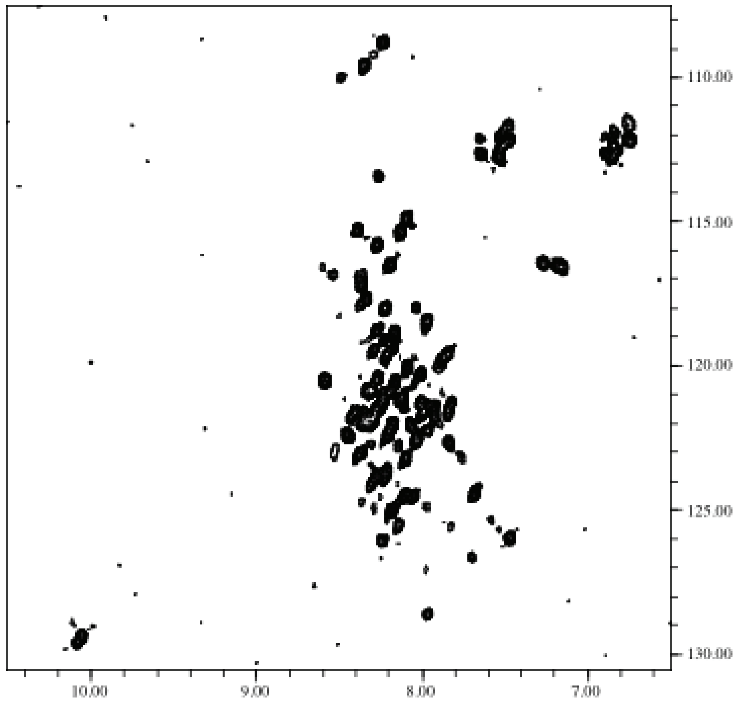


HSQC of Paip2 (1-75) in complex with RRM23

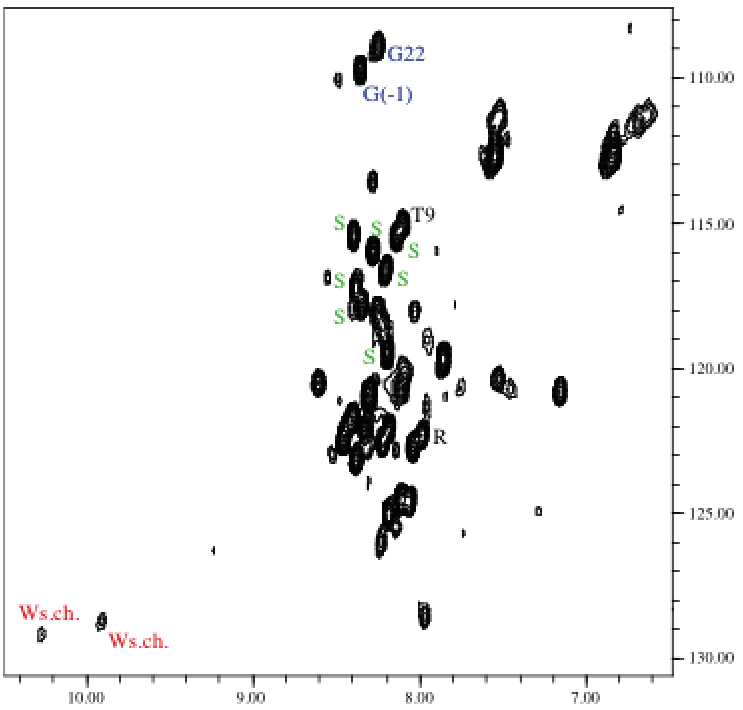

Fig. S4

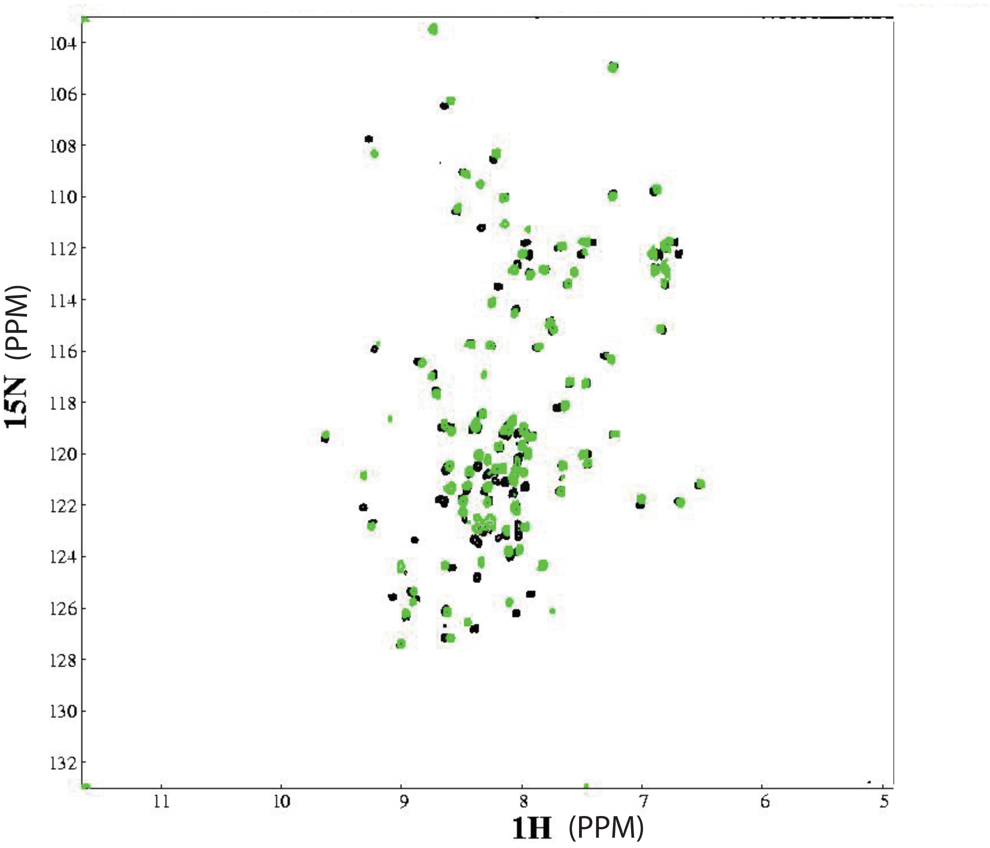

104
106
108
110
112
114
116
118
120
122
124
126
128
130
132
15N (PPM)
11
10
9
8
7
6
5
1H (PPM)

A

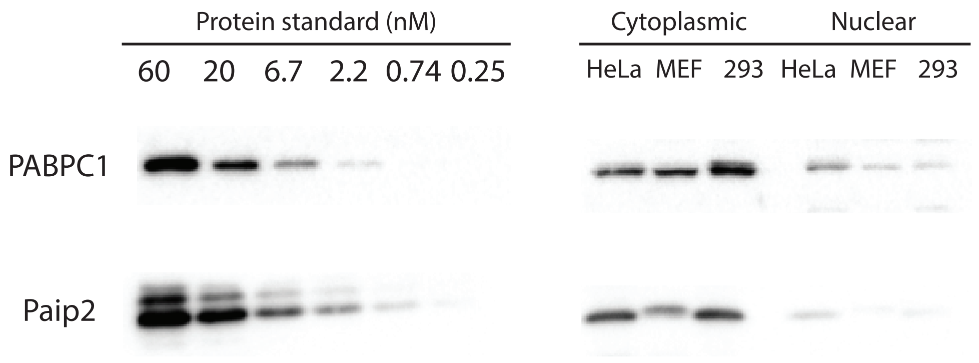


B

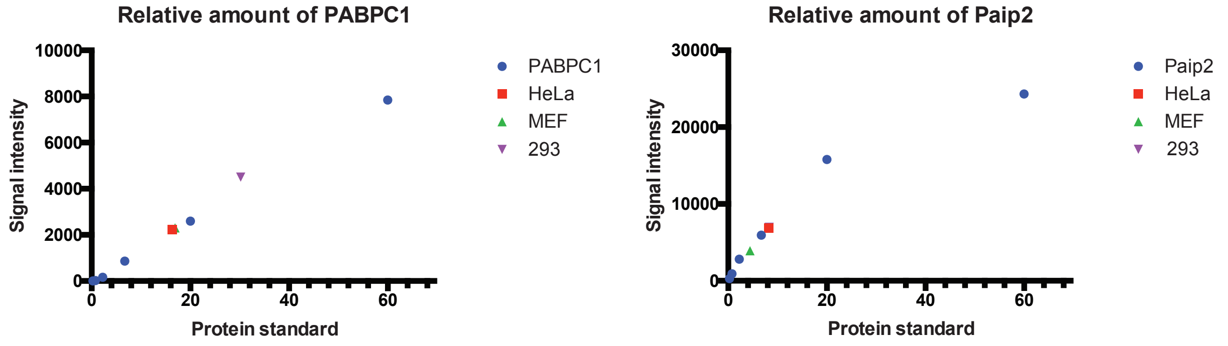


Relative amount of PABPC1 and Paip2 in cells (nM)

| | Hela CE | MEF CE | 293 CE | Hela NE | MEF NE | 293 NE |
|---|---|---|---|---|---|---|
| PABPC1 | 16.3 | 16.9 | 30.2 | 6 | 1.5 | 1 |
| Paip2 | 8.2 | 4.4 | 8.3 | 0.5 | -0.5 | -0.3 |
| PABPC1/Paip2 | 2 | 3.9 | 3.6 | - | - | - |

C

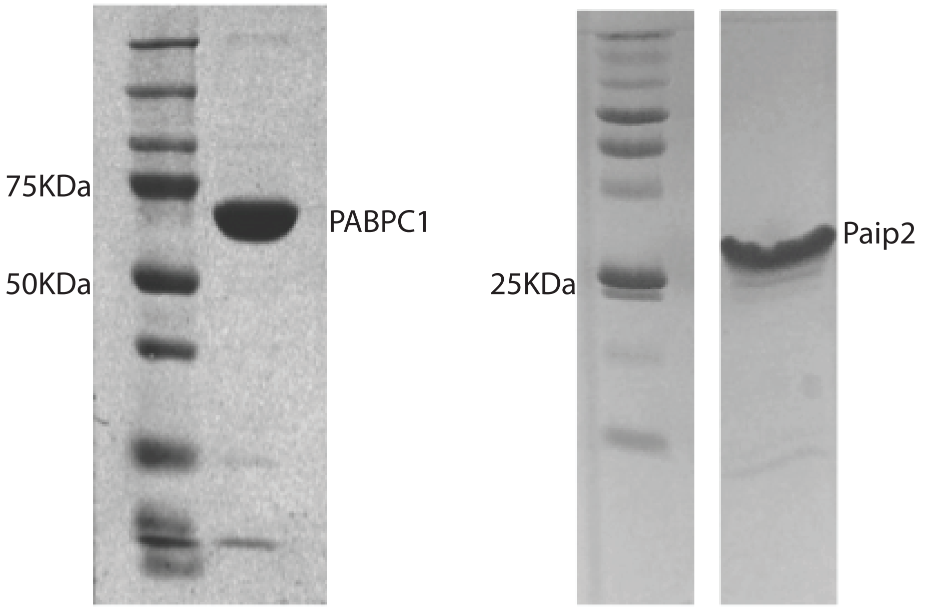
